# Introduced crops supplement rather than replace indigenous crops in an African center of agrobiodiversity

**DOI:** 10.1101/2022.03.30.486402

**Authors:** Chris Rampersad, Tesfu Geto, Tarekegn Samuel, Meseret Abebe, Marybel Soto Gomez, Samuel Pironon, Lucie Büchi, Jeremy Haggar, Jonathan Stocks, Philippa Ryan, Richard Buggs, Sebsebe Demissew, Paul Wilkin, Wendawek M. Abebe, James S. Borrell

## Abstract

- Crop diversity plays a major role in underpinning food security. It is especially important to smallholder and subsistence farmers, who often rely on crop diversity for stable and resilient production. Despite this, global expansion of a small pool of major crops and the associated homogenisation of global agricultural systems may decrease on-farm crop diversity.
- We surveyed 1,369 subsistence farms stratified across climate gradients in the Ethiopian Highlands, to characterise the richness and cultivated area of the 83 edible crops they contained. We further categorise these crops by their period of introduction to Ethiopia. We apply non-metric multidimensional scaling and mixed effects modelling to characterise agrisystem composition and test the impact of crop introductions.
- We find a significant positive relationship between introduced and indigenous crop richness, suggesting that crop introductions have tended to supplement rather than replace or reduce indigenous crop diversity. Geographically matched farms with higher proportions of introduced crops, had significantly higher overall crop richness. Analysis of socio-economic drivers indicates that both poverty and low accessibility are associated with reduced cultivation of modern introductions.
- We conclude that global patterns of major crop expansion do not necessarily result in agrobiodiversity loss for subsistence farmers, in our Ethiopian case study. Importantly, socioeconomic factors may strongly influence the farmers propensity to adopt novel species, suggesting targets for agricultural extension policies. Given the rapid climatic, economic and demographic changes impacting global food systems and the threats to food security these entail, robust indigenous agricultural systems supplemented with diverse introduced crops may enhance resilience.

## Introduction

Crop diversity plays a central role in delivering stable and resilient food production (Renard & Tilman, 2019). Among multiple benefits is evidence of increased productivity (Li *et al*., 2009), nutritional security (Nicholson *et al*., 2021), harvest asynchrony (Egli *et al*., 2020) and regulating ecosystem services (Kremen & Miles, 2012; Tamburini *et al*., 2020). Whilst crop diversity has been shown to have increased at the national scale during the Anthropocene, particularly via the Colombian exchange and green revolution (Khoury *et al*., 2016; Martin *et al*., 2019), the success of a small number of crops has also catalysed the homogenisation of global agrisystems (Khoury *et al*., 2014). This has led to concerns that the global expansion of these major crops could displace or replace indigenous crops (Shelef *et al*., 2017; Borrell *et al*., 2020; Khoury *et al*., 2022), with associated loss of long term local adaptation, indigenous knowledge and autochthonous resilience strategies (Seburanga, 2013; Raeboline *et al*., 2019).

Existing studies make extensive use of national crop production data from the United Nations Food and Agricultural Organisation (FAOSTAT 2021) to infer national crop diversity trends (Khoury *et al*., 2014; Martin *et al*., 2019; Mariani *et al*., 2021). However, to enable comparisons and consistency, crop data are often restricted to globally, or at least regionally important species and thus may overlook locally representative crops with a significant contribution to local agrobiodiversity and food security (Ulian *et al*., 2020). Furthermore, it is unclear how global trends in crop diversity are operating at the farm scale and the degree to which introduced crops are supplementing and integrating with, or replacing, local agricultural systems. Understanding the impact of crop introductions may inform strategies for enhanced food system stability and agrobiodiversity conservation, particularly under the pressures of poverty, population growth, and climate change (Borrell *et al*., 2020; Labeyrie *et al*., 2021).

Here, we investigate the impact of modern crop introductions to address knowledge gaps concerning local crop diversity trends at multiple scales, from farms to the agricultural systems that these comprise. We focus on the Ethiopian Highlands, a major Sub-Saharan African centre of crop diversity (Harlan, 1969), where over 80% of the population are engaged in subsistence agriculture. With a long history of crop domestication, Ethiopia represents a critical reservoir of agrobiodiversity and indigenous knowledge encompassing a range of major and minor crops suitable for investigating local trends in crop diversity and agricultural homogenisation. We propose that interactions between introduced and indigenous crops are mediated by three possible responses: 1) *Supplementation*, where farmers integrate novel crops with desirable traits into existing agricultural systems. 2) *Replacement*, where an introduced crop with a desirable trait is favoured over an existing species. 3) *Reduction*, where farmers rely on fewer introduced crops to meet their needs by replacing pre-existing species (Figure 1). We also highlight that changes in diversity emerging from crop introductions may be rapid or require a protracted period to approach an equilibrium.

**Figure 1.**
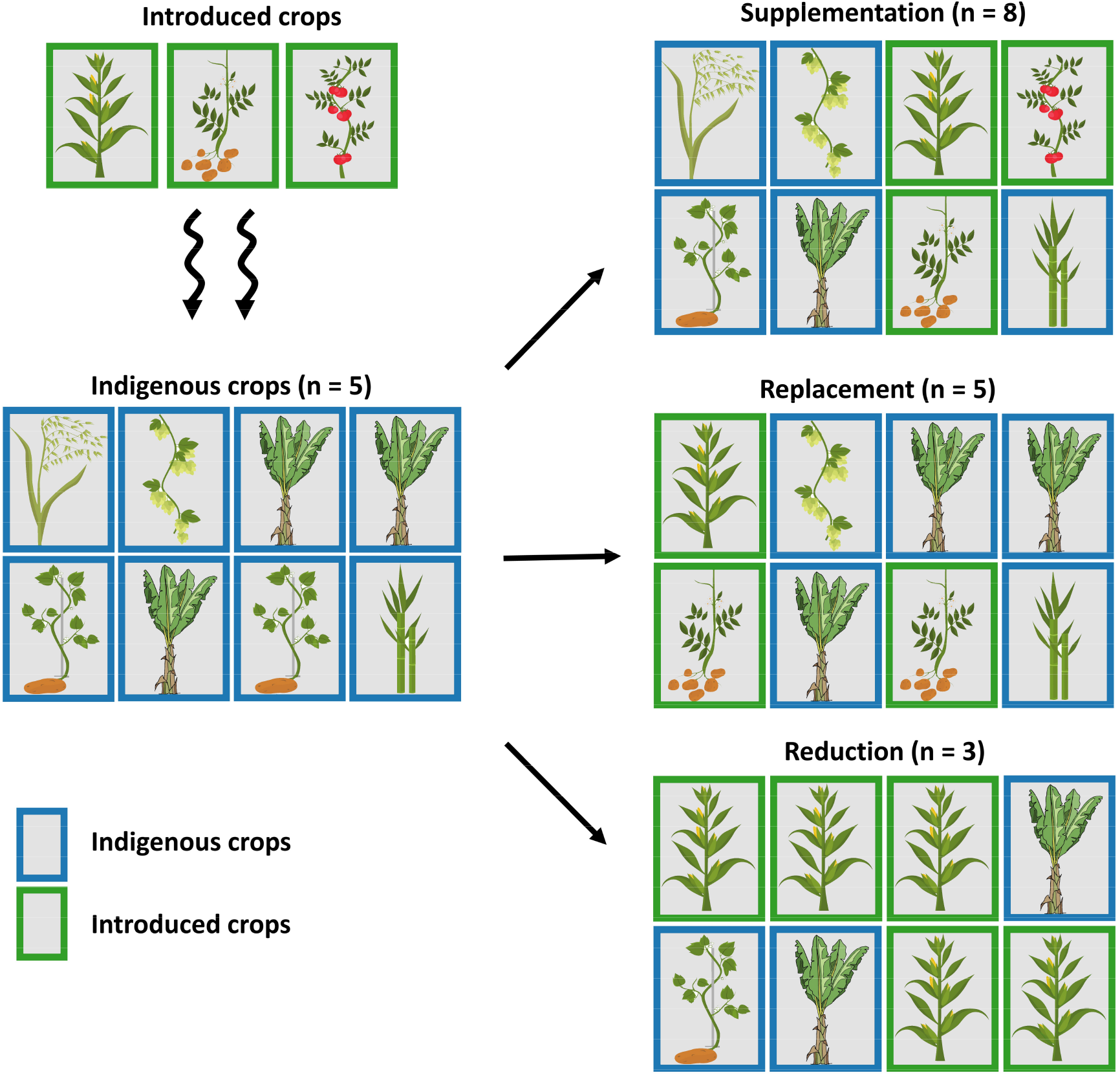
Hypothesised impact of crop introductions (Modern/Early-modern) on pre-existing (Indigenous/Historic) crop diversity. Reduction of crop diversity occurs where multiple indigenous crops are substituted by fewer species of introduced crops, reducing overall crop richness. Replacement involves 1:1 exchange of indigenous for introduced crops, with overall richness remaining constant. Supplementation is where additional introduced crops are integrated into the existing system, increasing overall crop richness. Multiple mechanisms could operate concurrently at farm and landscape scales.

Using a novel dataset of comprehensive farm surveys across Southwestern Ethiopia, combined with information on the period of introduction for 83 crops, we frame our analysis with three questions. First, to what extent have introduced crops from across multiple time periods become integrated within Ethiopian agricultural systems? Second, what socio-economic drivers influence the proportion of introduced crops cultivated on farms? Third, do introduced crops supplement, replace or reduce indigenous crop diversity, and what is the impact at farm and agricultural system scales? Finally, we consider how these findings can inform future agrobiodiversity management.

## Methods

### Stratified farm surveys

We sampled 1,369 farms across eight transects in Southwestern Ethiopia between February and July 2019. Transects were orientated perpendicular to elevational and environmental gradients ensuring representative coverage of regional agroecological variation. Transects were 20-50 km in length and sampling ranged from 1,200-3,240m asl (Figure S1). In each farm, we recorded the presence and cultivated area of all actively managed plant species. For some perennial crops that tend to be grown in low numbers outside of farmer fields, the number of individuals was recorded and converted using a constant area per individual, derived from an initial measurement of >10 individuals per crop (e.g. 0.002 ha for avocado trees). While 124 crop species were recorded in total, for subsequent analyses we retained only those that yield human edible products (n = 83, Table S1); excluded species were predominantly used for timber and fodder. Each farm’s total harvest area was calculated by summing the area of all crops cultivated. We also recorded overall farm area by integrating the area of wood lots (mostly eucalyptus), pasture and other non-cultivated land.

### Characterisation of crop introduction categories

To investigate interactions between indigenous and introduced crops, we sought to determine the period of introduction of the 83 crops in our dataset. We note that our analysis focuses on crop species, and does not account for landrace or varietal diversity (see Discussion). We surveyed the literature and assigned crops to one of four categories: i) Indigenous, comprising crops for which there is evidence of domestication, or the occurrence of wild progenitors in Ethiopia (n=25). ii) Historic, crops likely introduced before c. 1500 AD, often with evidence of a secondary centre of diversity within Ethiopia (n=25). iii) Early modern, those introduced during a period of increased international exposure from 1500-1900 arising in part from the Colombian Exchange (Williams, 2017) (n = 18). iv) Late modern, those introduced in the last century, predominantly associated with global trade and the green revolution (n=15). We used analysis of variance (ANOVA) to test for differences in the frequency and cultivated area among the four crop groups. For some subsequent analyses we group indigenous and historic versus early and late modern crop classes. All analyses were performed in R v3.6.3 (R Core team 2019).

### Agricultural system composition and integration of modern crops

To understand whether on-farm crop assemblages are partitioned into distinct agricultural systems in Ethiopia, we applied non-metric multidimensional scaling (NMDS) based on a scaled Bray-Curtis dissimilarity matrix. NMDS was implemented in the R package vegan (Oksanen *et al*., 2019), over two to six axes, with up to 500 random starts and 2000 iterations. We treated farms as observation units and crop species as descriptors. We used the *envfit* function to determine the extent to which crop species covaried with each axis, and applied analysis of variance to test for differences in correlation by period of introduction. We plot NMDS values with the corresponding Ethiopian agroecological zones by calculating 95% ellipses around farms in each elevational band. These zones have traditionally been used to describe distinct systems based on elevation as a proxy for climate (Hurni, 1998).

To further evaluate the relative importance of introduced crops to agrisystem composition we investigated patterns of crop co-occurrence, using network visualisation in the iGraph package (Csardi & Nepusz, 2006). We weighted node size by frequency of crop occurrence across farms and plotted undirected edges proportional to the number of farms in which the species co-occur. We then used analysis of variance to compare eigenvector centrality by crop category (i.e. indigenous, historic, early modern or late modern), which measures the influence of a node in a network. Higher scores denote nodes that are themselves connected to many other high scoring nodes, an indication of crop importance within the regional agricultural system.

### Socioeconomic drivers of introduced crop cultivation

We used mixed effects models implemented in the R package nlme (Pinheiro et al, 2020) to investigate the socioeconomic drivers influencing the proportion of (early and late) modern crops cultivated on farms. We used both species richness and cultivated area as response variables. Fixed effects comprised total harvested area (ha) and number of livestock, both serving as proxies for farmer affluence. We also applied a proxy for regional affluence, comprising travel time to the nearest town (extracted at 1km resolution from Ethiogis 3; www.ethiogis-mapserver.org) and a compositive poverty index. The latter index is a composite of six regional health, wellbeing and development indicators including access to safe drinking water, months of food insecurity, child vaccination, child stunting and male and female literacy rates, all derived from the Ethiopia 2016 Demographic and Health Survey (USAID, 2021) (see Supplementary Information). The eight transects were treated as a random effect and a Gaussian covariance structure was applied to farm coordinates to account for spatial autocorrelation. Models were evaluated using AIC and examination of residual plots following Zuur et al. (2009).

### Testing whether introduced crops supplement, replace or reduce agrobiodiversity

We applied three approaches to test the impact of introduced crops at both farm and agricultural system scales. First, we used a linear mixed effects model to evaluate the relationship between counts of indigenous versus modern introduced crops within farms, while accounting for spatial autocorrelation using a Gaussian correlation structure with transect as a random effect. We examine this relationship against our hypotheses of crop supplementation, replacement or reduction by farmers (Figure 1). Second, we tested the impact of the proportion of farm area allocated to modern crops on overall farm crop richness and evenness using Simpson’s diversity index (Hurlbert 1971). To do this we applied similarly structured linear mixed effects models, including a quadratic fixed variable to accommodate a non-linear relationship in the response.

Third, to assess the impact at the agricultural system scale, we collated our dataset of 1,369 farms into clusters of ten farms, based on geographic proximity. From each cluster we then selected the farm with the highest and lowest proportion of modern crops (considering early and late introductions jointly) which generated 137 pairs. This approach aims to maximise the chance of detecting an effect of modern crop introductions, whilst controlling for the effects of bioclimatic and cultural variables, as farms from the same locale are expected to experience similar conditions. We applied our previous NMDS model to visualise changes in overall crop richness between these two groups of high versus low diversity of modern crops. Following an approach similar to Khoury et al. (2014) we fitted ellipses to each set of farms and compared the volume of multivariate space encompassed by each group. We tested for differences in the composition and dispersion of these groups using analysis of similarities (Oksanen *et al*., 2019), which compares multivariate rank order of dissimilarity values.

## Results

### Agricultural system composition and integration of new crops

Our analysis of crop introductions finds that Ethiopia has been characterised by the continuous arrival of novel crop species over several millennia (Figure 2). Despite this, the largest proportion of crops recorded in our study area were of indigenous (n = 25, 30.1%) or historic (n = 25, 30.1%) origin, consistent with Ethiopia’s role as a major center of crop domestication as well as an ancient secondary center for crops such as wheat (Table S1). We found no significant difference in the frequency of on-farm occurrence (F_3,79_=0.54, p=0.65), or cultivated area (F_3,79_=0.77, p=0.51) among the four categories of crop origin (Figure 2). Fruit crops, followed by roots and vegetables represent the most frequent (successfully) introduced crop types of the modern era.

**Figure 2.**
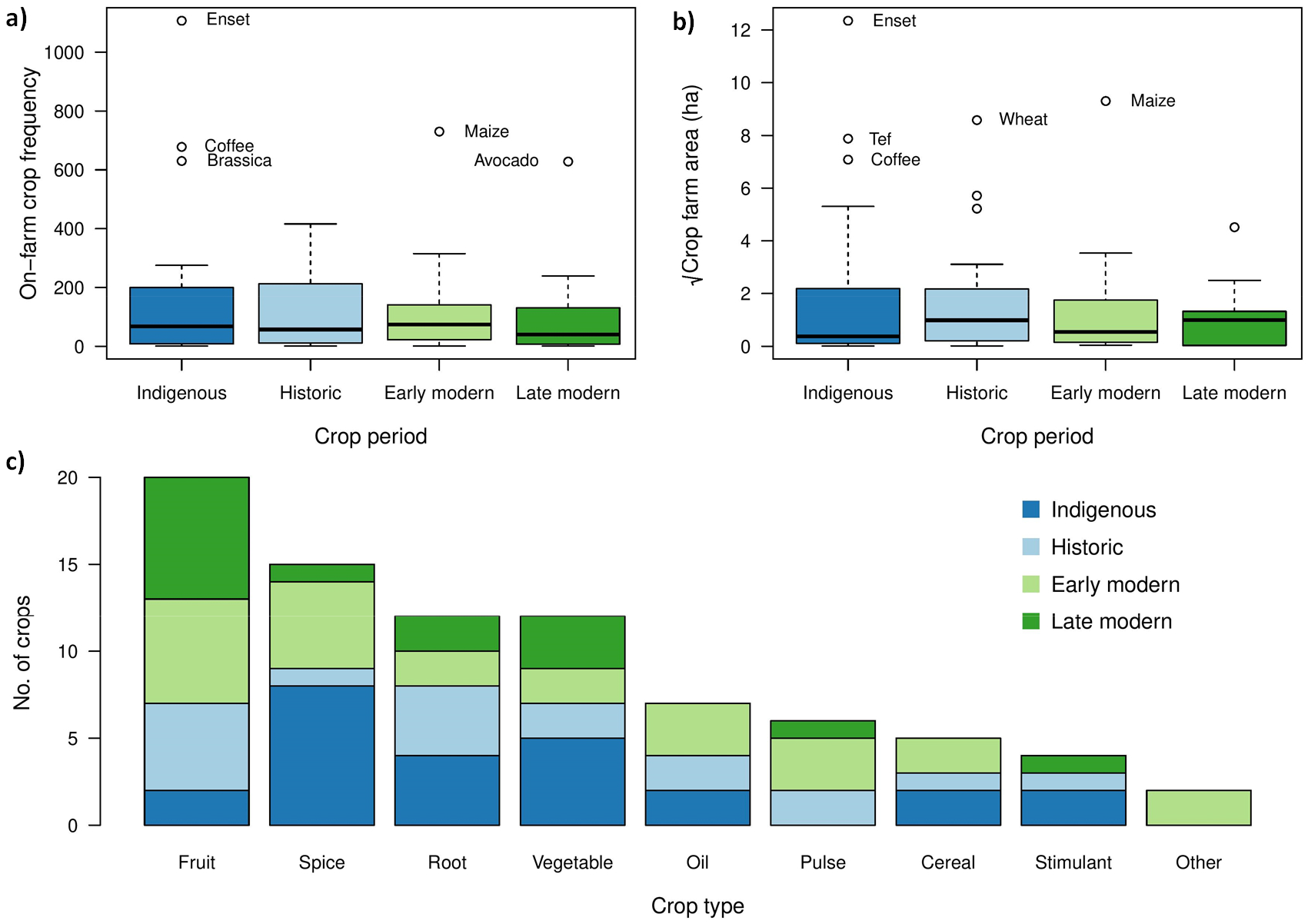
Crop frequency, area and period of introduction across 1369 surveyed farms in the Ethiopian highlands. a) Boxplot of observation frequency for indigenous and introduced crops, grouped by period of introduction. b) Boxplot of total cultivated farm area for crops by period of introduction. c) Stacked bar chart of crop types occurring in the study area; category ‘Other’ comprises olive (*Olea europaea*) and sugarcane (*Saccharum officinarum*).

Farmers cultivate up to 29 edible species (mean = 8.08, SD = 3.49) in the study area and total harvest area varies from 0.004 - 3.07 ha (mean = 0.48 ha, SD = 0.41 ha). We found no evidence of clustering in crop assemblages in a fitted a four-dimensional NMDS (stress = 0.12; Figure 3). Similarly, crop assemblages in Ethiopian Dega and Weyna Dega zones strongly overlapped. Crops that covaried most strongly with NMDS axes encompass both indigenous and recently introduced species, including wheat (R_2_ = 0.44), maize (R_2_ = 0.37), tef (R_2_ = 0.35) and enset (R_2_ =0.25). Overall, we found no significant difference in correlation across species with different periods of introduction (F_3,79_ = 0.9, p = 0.45). In other words, no group of crops disproportionately contributed to variation or structure in the regional agricultural system. Similarly, network analysis identified no evidence of clustering in on-farm crop assemblages (Figure 4). The most frequently recorded and interconnected crops included indigenous (enset, coffee) and introduced (avocado, maize) representatives. We found no significant difference in eigenvector centrality across indigenous and introduced crops from different periods (F_3,79_ = 0.45, p = 0.72), suggesting that variation in the connectedness of nodes (i.e. crops) is not associated with the length of time since introduction.

**Figure 3.**
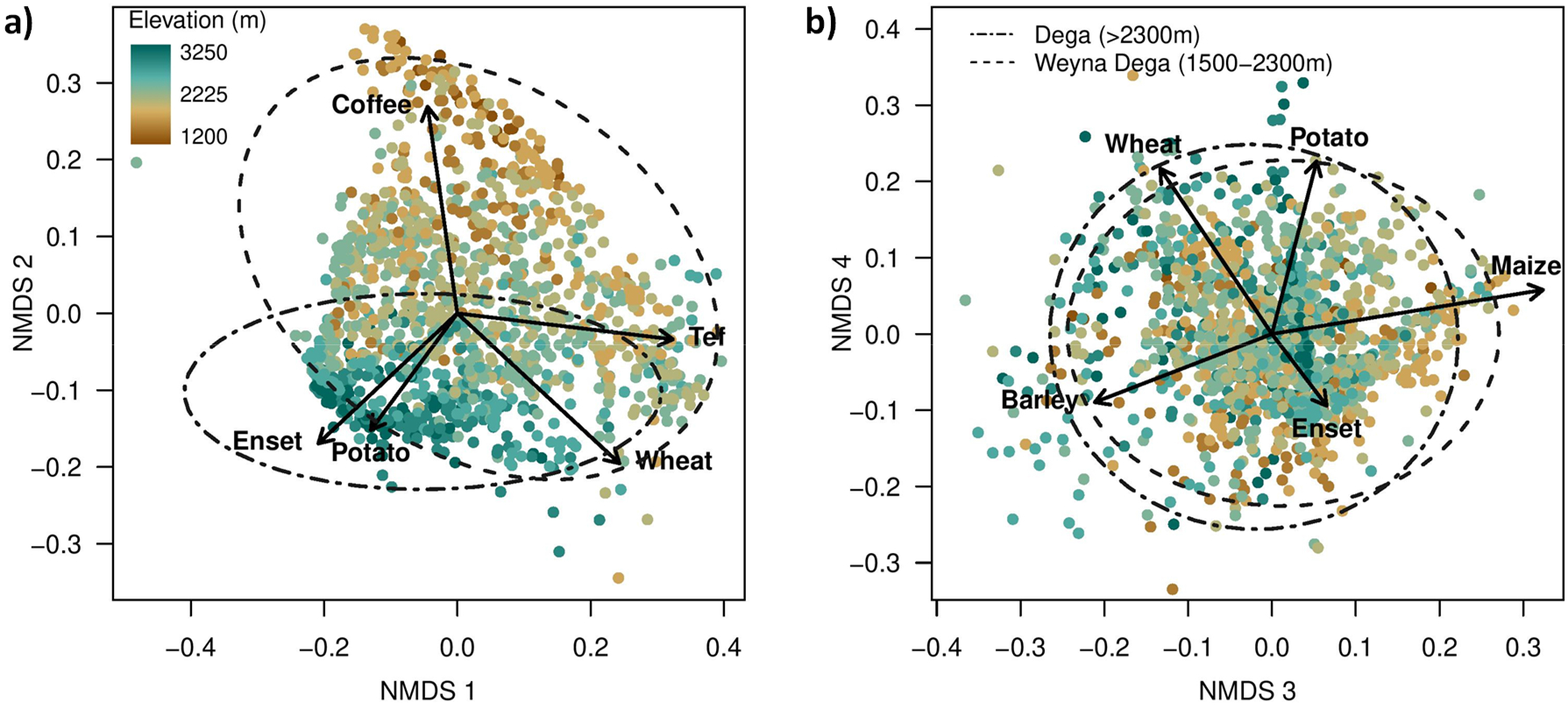
Non-metric multi-dimensional scaling analysis (NMDS) of farm crop composition in the Ethiopian highlands. Each point is one of 1,369 farms, with plots showing a) the first and second axes, and b) the third and fourth axes. Ellipses encompass the environmental space occupied by farms in two predominant Ethiopian agroecological zones, Dega and Weyna Dega, at 95% confidence. Arrows indicate direction and magnitude of correlation for key species on the ordination.

**Figure 4.**
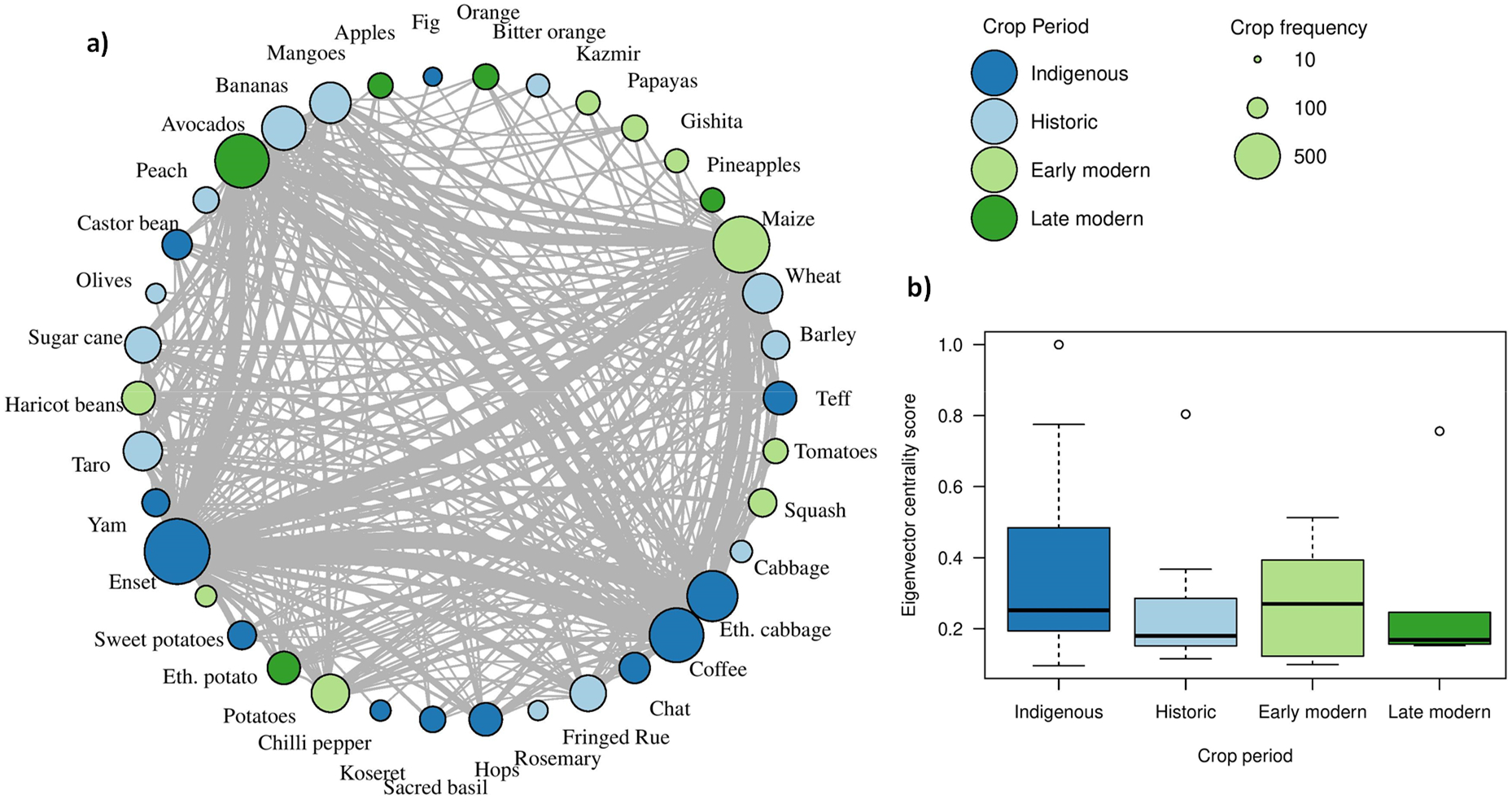
Agrisystem network composition and eigenvector centrality analysis. a) Network analysis where node size denotes the number of farms with crop presence and edge width between nodes corresponds to frequency of co-occurrence. Crops occurring on less than 5% of farms are omitted for plotting purposes. b) Boxplot of eigenvector centrality across indigenous and introduced crops (grouped by crop period).

### What influences the proportion of modern introduced crops?

Mixed effects models identified no significant variables that influenced the proportion of modern crop introductions found on farms, though increasing travel time had a marginally non-significant negative effect (Table 1). However the proportion of cultivated area allocated to crop introductions was strongly negatively associated with poverty. In addition, increasing travel time and total cultivated area were also associated with lower proportion of introduced crop cultivated area. In contrast, head of livestock (a proxy for farmer wealth) was associated with a greater proportion of introduced crop area.

**Table 1.** Influence of socio-economic variables on the proportion of modern introduced crop richness (count) and area cultivated (ha).

| Variable | Proportion of modern introduced crops |  |  |  |
| --- | --- | --- | --- | --- |
|  | Richness (count) |  | Area (ha) |  |
|  | t-value | p-value | t-value | p-value |
| Total cultivated area | -0.97 | 0.33 | -2.75 | 0.01 |
| Travel time to main town | -1.84 | 0.07 | -3.27 | <0.01 |
| Total Livestock | -0.13 | 0.9 | 3.03 | <0.01 |
| Poverty composite variable | -1.36 | 0.17 | -6.24 | <0.01 |

### Do modern introduced crops supplement, replace or reduce existing crop diversity?

At the farm scale we find a significant positive relationship between indigenous and introduced crop richness (t_1360_ = 12.77, p < 0.001) (Figure 5). In comparing this with our hypotheses (Figure 1), it suggests that introduced crops have predominantly supplemented, rather than replaced or reduced existing indigenous crop diversity. Consistent with this, we find significant concave relationships between the proportion of introduced crops and both overall crop richness (X, t_1358_ = 6.88, p < 0.001; X^2^, t_1358_ = −12.06, p < 0.001) and Simpson’s evenness index (X, t_1358_ = 10.54, p < 0.001; X^2^, t_1358_ = - 21.68, p < 0.001). This shows that higher overall crop richness and evenness on farms is associated with intermediate proportions of both indigenous and introduced crops (Figure 6). We note that the highest overall richness is observed at a proportion of 40% introduced crops, but that 81% of the surveyed farms are below this value.

**Figure 5.**
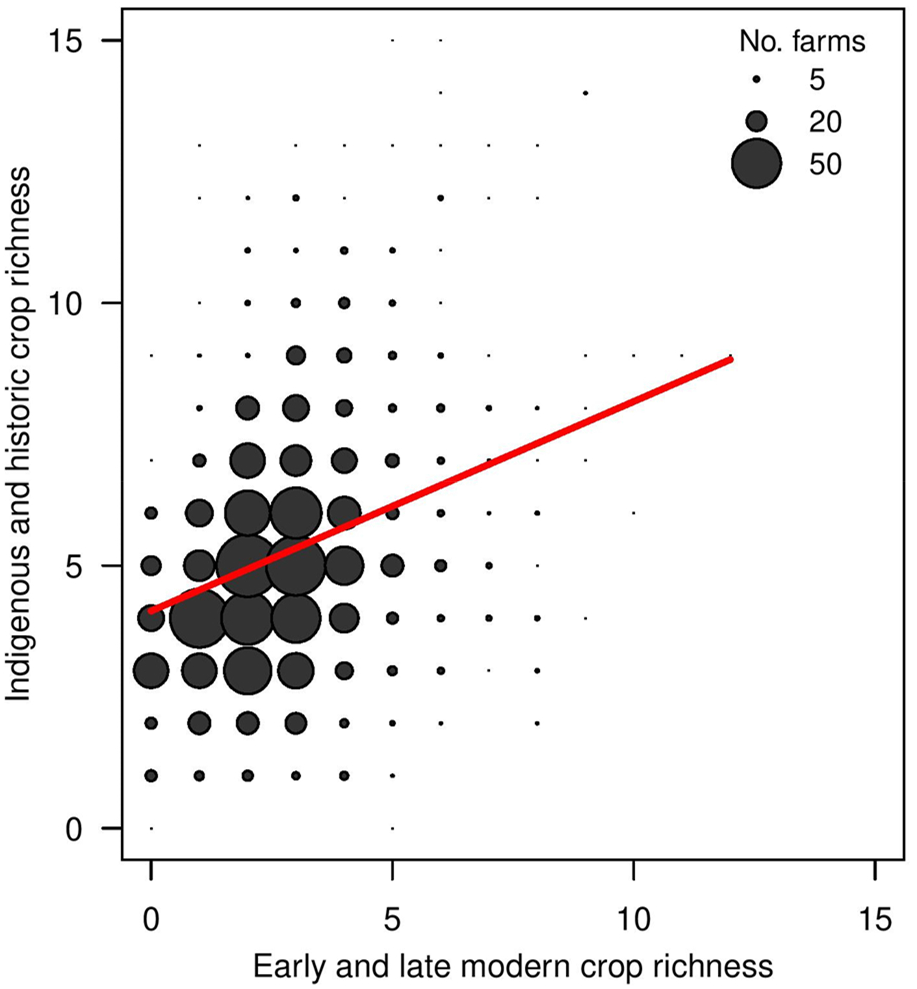
Relationship between early and late modern crop richness versus indigenous and historic crop richness across 1369 farms in the Ethiopian Highlands. The red line indicates the fit of a linear mixed effect model and points are scaled by number of farms.

**Figure 6.**
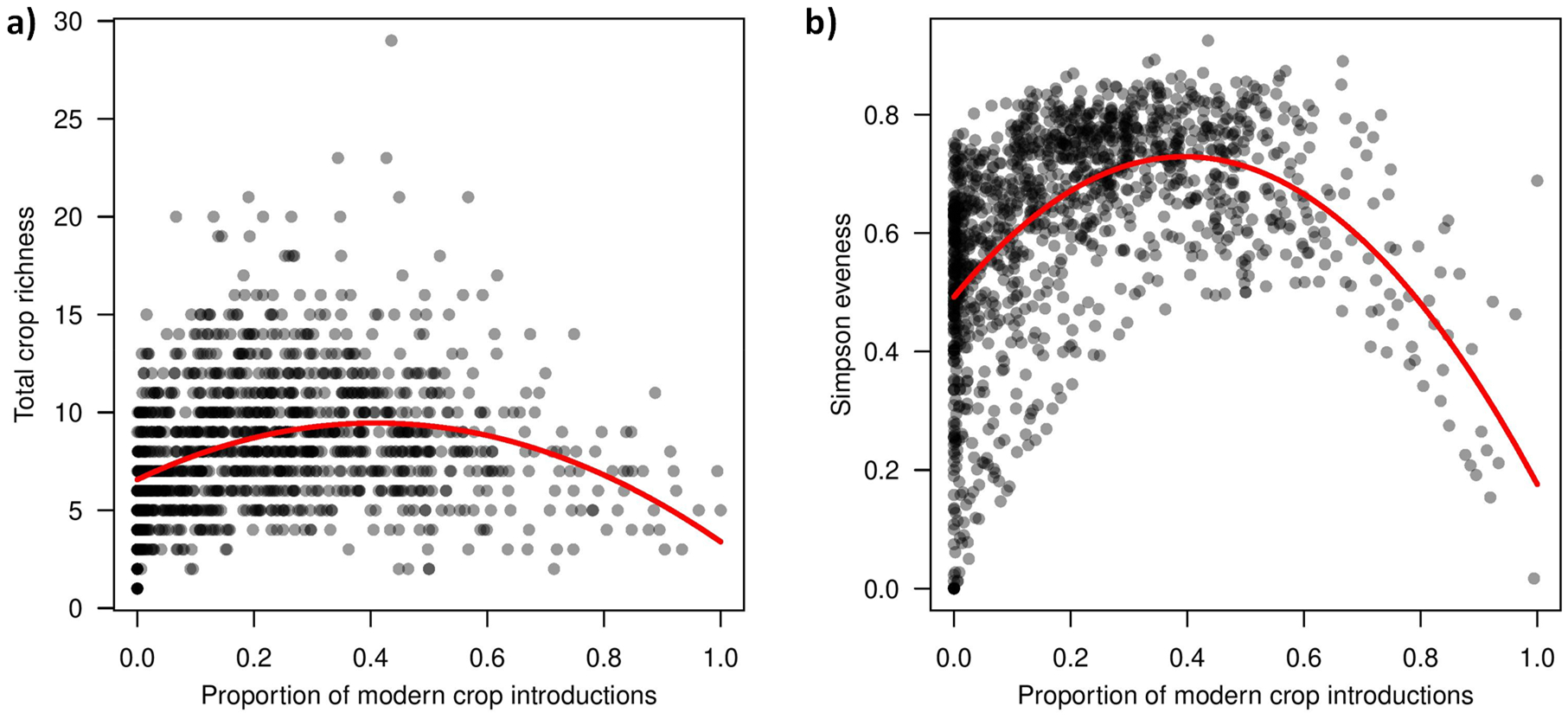
Impact of modern crop introductions on overall crop richness and evenness. Plots show associations between the cultivated proportion of modern crop introductions versus a) overall crop richness on farms and b) Simpson’s evenness index. Trend lines, shown in red, were fitted using linear mixed effects models.

At the agricultural system scale, we used geographic proximity to match and compare a subset of farms containing a high proportion of recently introduced crops (mean proportion modern = 0.48), with a subset of farms containing a low proportion of introduced crops (mean proportion modern = 0.04) (Figure 7). Despite maximising the possible difference in the proportion of introduced crops across these two subsets, we found that this only significantly explained a very small amount of variation (R_2_ = 0.08, p = 0.001). Further examination suggests that this difference may be driven by higher variance, with the ‘high proportion’ group occupying a 12% larger volume in multivariate space, indicating greater diversity in crop species and relative crop area composition across the agricultural system.

**Figure 7.**
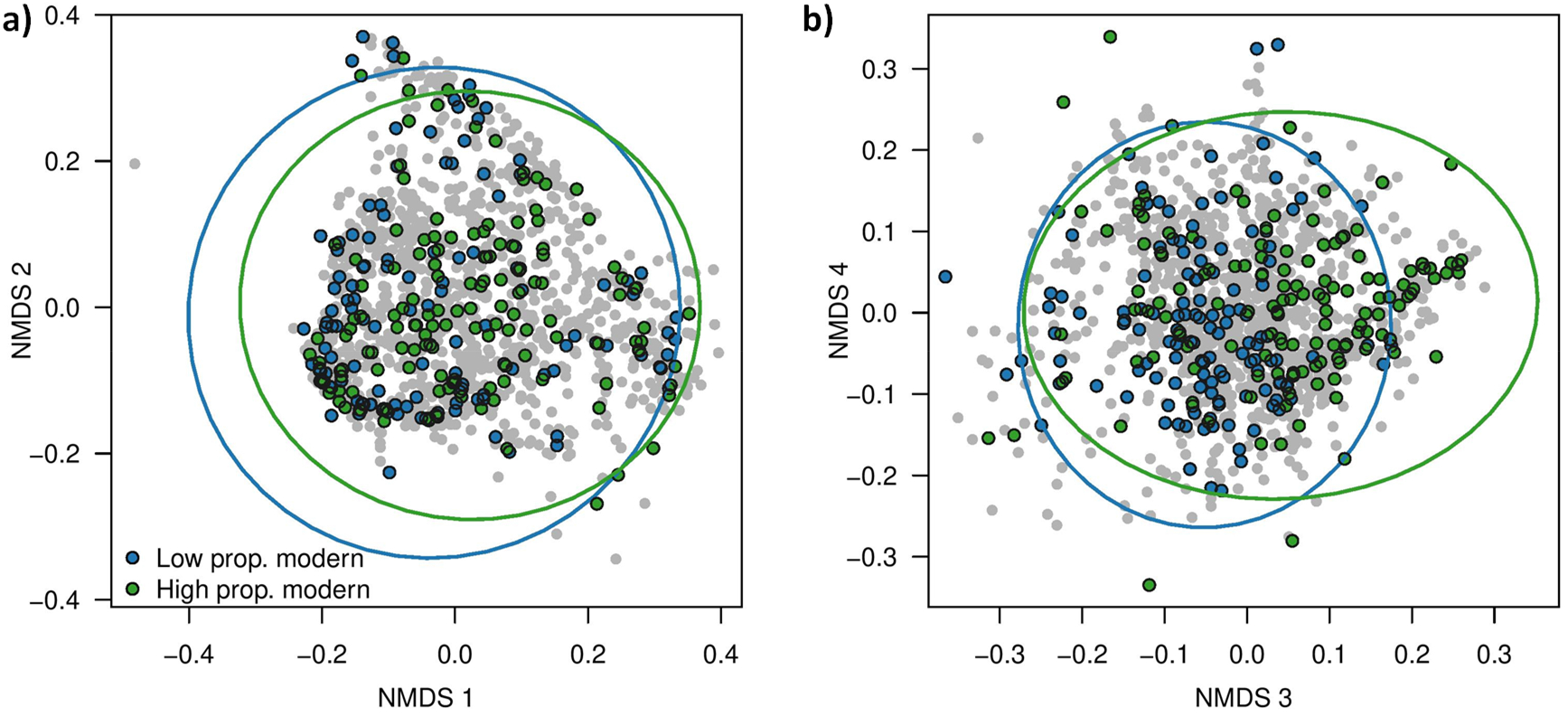
Impact of crop introductions on agrisystem scale diversity. NMDS of all farms with coloured points indicating two groups of climatically matched farms (n=137 each) comprising low and high proportions of modern crop introductions. Ellipses capture 95% of multivariate space for each group.

## Discussion

Global loss of indigenous crop diversity as a result of introduced crop expansion is a major food security concern and could undermine the sustainability of food systems (Renard & Tilman, 2019). Here, we provide evidence that introduced crops tend to supplement rather than replace or reduce indigenous crop diversity in Ethiopia. This is supported by several lines of evidence. First, we observe a positive relationship between the number of modern crop introductions present on a farm and overall crop diversity (Figure 5). This suggests that farmers are integrating novel crops, without reducing their existing suite of species. Second, we observe that farms with intermediate proportions of modern crop introductions show the highest overall crop richness and evenness (Figure 6). This indicates that in most farms, crop introductions do not substantially reduce the cultivated area of indigenous crops. Additionally, more extensive cultivation of introduced crops may increase overall crop richness and evenness in most surveyed farms, as ∼80% of these are to the left of our models’ apex. Thirdly, using a matched pairs approach, whereby farms have similar geography and climate, but a 10-fold difference in the proportion of introduced crops, we report a 12% expansion of multivariate space that characterises agricultural system diversity. Taken together these three approaches provide evidence to support our supplementation hypothesis in which modern crop introductions are contributing additional agrobiodiversity to Ethiopian agrisystems while not having a deleterious effect on indigenous species richness.

A second key finding is that introduced crops appear to be rapidly integrated into indigenous agricultural systems. For example, NMDS across surveyed farms found little evidence of clustering of crop compositions (Figure 3), contrary to previous reports that southwestern Ethiopian agrisystems are differentiated (Abebe *et al*., 2010). Within this system we found that indigenous and introduced crops form a highly network (Figure 4a). For example, both enset (an indigenous starch staple, cultivated only in South-western Ethiopia) and avocado (a recently introduced fruit) occur frequently with a wide variety of other crops. We found no significant difference in connectedness between groups of indigenous or introduced crops regardless of the timing of introduction (Figure 4b). This is surprising because we might expect more recent introductions to be less well integrated (Ali & Erenstein, 2017; Uduji & Okolo-Obasi, 2018), suggesting that subsistence farmers may integrate new crops in a relatively short period.

In assessing the socio-economic drivers of the proportion of introduced crops found on surveyed farms, we identified poverty as a major negative driver. We also detected negative associations with total cultivated area on farms, and accessibility (Table 1). Conversely, the head of livestock – a proxy for wealth – was positively associated with a larger proportion of introduced crops in surveyed farms. This suggests that access to modern introduced crops is associated with development, and importantly can be targeted by appropriately designed rural development policies (Welteji, 2018). Additionally, modern introduced crops may be more profitable to sell, and so have higher uptake among more commercially oriented farmers, who are likely those with higher access to markets, and sufficient wealth to meet their subsistence needs.

Previous studies have reported a global narrowing of crop species that contribute to world food supplies (Khoury *et al*., 2014; Martin *et al*., 2019). While expansion of major crops has resulted in relatively high species richness nationally, this has been associated with increasing similarity of species composition internationally. Our farm-scale findings for Ethiopia complement and contrast these analyses to provide a clearer assessment of ongoing shifts in agrisystem composition. At the local scale we find that farms cultivating more introduced crops have a higher number of crop species overall (Figure 5), higher evenness across species (Figure 6) and result in agricultural systems with more variation in crop composition (Figure 7). We suggest that the understandable omission of numerous relatively underutilised or locally important species, particularly from smallholder subsistence farms, underestimates farm-scale agrobiodiversity in global agrobiodiversity datasets such as FAOSTAT (2021). For example, 36% of species in this study are not included in the global analysis of Martin et al. (2019), including major regional staples such as enset (*Ensete ventricosum*) (Borrell *et al*., 2019). Similarly, while we lack time-series data at the local scale and do not assume that national agricultural systems have reached an equilibrium, we also do not see trends in crop importance associated with period of introduction. In other words, the length of time a species has been in Ethiopia, does not appear to be associated with its relative importance (Figures 2, 4). These findings support a growing body of work that modern agricultural transformations have not invariably led to large scale agrobiodiversity loss (Renard *et al*., 2016; Khoury *et al*., 2022).

We note several caveats and limitations of our study. Foremost is the widely reported evidence of loss of traditional landraces in favour of improved and often introduced genotypes (Thormann and Engels 2015), and the associated loss of numerous generations of locally adaptive evolution (Kassahun *et al*., 2021). Our analysis does not capture changes at the intraspecific level, and thus we have not quantified landrace replacement though it is likely occurring. Second, we note that dating and characterising the period of introduction for many crops is challenging, particularly where region-specific archaeobotanical or historical evidence is lacking. Nevertheless, we have attempted to draw justifiable inferences based on records of crop expansion, genetic studies and known cultural contact (e.g. the Colombian Exchange), and our results are robust to modest adjustments in the categorisation of crop introduction periods (see Supplementary Table S1). Our study captures a contemporary time period, so while we detect no loss of indigenous crop diversity, we cannot rule out the possibility of a protracted decline in the future towards equilibrium (conceptually similar to ‘Extinction Debt’, see Kuussaari et al. 2009), or an unrecognised ancient decline. Finally, we note that as a globally important center of crop diversity and domestication, with a regionally unique climate, Ethiopia may not be widely representative of trends in other countries or regions. Indeed, the stability of indigenous agrisystems in our study may be partly due to the comparatively minimal extent of colonial era agri-polices, partly shielding Ethiopia from the effects of mid-20^th^ century green-revolution influenced interventions (Till, 2021).

In the future, emerging climate and demographic pressures may require the relatively rapid shift and rearrangement of current crop distributions (Sloat *et al*., 2020; Koch *et al*., 2022), as well as adoption of new and better adapted species and agrisystems (Pironon *et al*., 2019; Borrell *et al*., 2020). In Ethiopia, the long-term maintenance of crop diversity over multiple periods of origin, combined with rapid integration of novel species suggests that these agrisystems may be relatively adaptable and resilient. Future research should aim to bridge the gap between farm and national scale agrobiodiversity trends.

## Acknowledgements

This work was supported by the GCRF Agrisystems award entitled, ‘Landscape scale genomic-environment data to enhance the food security of Ethiopian agri-systems’ [Grant No. BB/S014896/1]. JB and SP were additionally supported by Future Leader Fellowships at the Royal Botanic Gardens, Kew.

## Author contributions

C.R. and J.B. designed the study, with substantial input from all authors; C.R., T.G., T.S., M.A., W.A. and J.S. collected data; C.R. and J.B. analysed the data with help from M.S.G., S.P. and L.B.; J.B., J.H., R.N., S.D., W.A. and P.W. secured funding. J.B. and W.A. provided supervision. C.R. and J.B. wrote the manuscript. All authors contributed to the final version.

## Research ethics

All participant farmers gave prior informed consent to participate in this study with the principals and aims explained in the predominant local language or Amharic. The resolution of identifiable information such as coordinates has been aggregated.

## Data availability

Databases of crop origin and anonymised farm composition are provided in Supplementary tables S1 and S2.

