## Supplementary Materials for "Introduced crops supplement rather than replace indigenous crops in an African center of agrobiodiversity"

### Composite poverty index

The poverty index is a composite of six regional health, wellbeing and development indicators derived from the USAID Demographic and Health Surveys Programme. These include percentage of children aged 12-23 months who had received all 8 basic vaccinations (USAID, 2016a); percentage of children under age five years stunted (USAID, 2016b); percentage of men age 15−49 who are literate (USAID, 2016c); percentage of women age 15−49 who are literate (USAID, 2016d); percentage of currently married or in union women with an unmet need for family planning (USAID, 2016e) and percentage of the population living in households whose main source of drinking water is an improved source (USAID, 2016f). We scaled all variables so that greater values were indicative of more severe health or development deficits and ranged from 0-1, and averaged all variables to derive a 5km scale composite for Ethiopia.

### Tables

**Table S1.** Crop genera and species identified across 1369 farm surveys in Southwestern Ethiopia, together with putative origin and crop type following categories of the Central Statistics Agency of Ethiopia.

| **Genus** | **Species** | **English name** | **Amharic name** | **Period** | **Type** | **References** |
| --- | --- | --- | --- | --- | --- | --- |
| *Eragrostis* | *tef* | Teff | Teff | Indigenous | Cereal | (Harlan, 1969; Zeven & Zhukovsky, 1975) |
| *Hordeum* | *vulgare* | Barley | Gebs | Historic | Cereal | (Brush, 2000; Phillipson, 2012) |
| *Sorghum* | *bicolor* | Sorghum | mashla | Indigenous | Cereal | (Harlan, 1969) |
| *Triticum* | *aestivum* | Wheat | sindea | Historic | Cereal | (McCann, 1995; Phillipson, 2012) |
| *Zea* | *mays* | Maize | bekolo | Early modern | Cereal | (Abate *et al.*, 2015) |
| *Ananas* | *sp.* | Pineapples | Ananas | Late modern | Fruit | (Gessesse *et al.*, 2019) |
| *Annona* | *sp.* | Gishita | Gishta | Early modern | Fruit | (Lawal, 2014) |
| *Artocarpus* | *heterophyllus* | Jackfruit |  | Early modern | Fruit | (d’Eeckenbrugge *et al.*, 2019) |
| *Carica* | *papaya* | Papayas | Papaya | Early modern | Fruit | (Harlan, 1969) |
| *Casimiroa* | *edulis* | Kazmir | Kazamora | Early modern | Fruit | (Satheesh, 2015; d’Eeckenbrugge *et al.*, 2019) |
| *Citrus* | *aurantium* | Bitter orange | Lomi | Historic | Fruit | (d’Eeckenbrugge *et al.*, 2019) |
| *Citrus* | *sinensis* | Orange | Komtatie | Late modern | Fruit | (Engels et al., 1991) |
| *Citrus* | *aurantifolia* | Lemon/lime | Trngo | Historic | Fruit | (Goettsch, 1991; d’Eeckenbrugge *et al.*, 2019) |
| *Citrus* | *medica* | citron | Brtukan | Historic | Fruit | (d’Eeckenbrugge *et al.*, 2019) |
| *Ficus* | *sp.* | Fig | Warka/Shola | Indigenous | Fruit | (Plants of the World Online \| Kew Science 2021) |
| *Fragaria* | *x ananassa* | Strawberry | Yemdir -'njorie | Late modern | Fruit | (Etissa *et al.*, 2016) |
| *Malus* | *sp.* | Apples | Pom | Late modern | Fruit | (Melke & Fetene, 2014) |
| *Mangifera* | *sp.* | Mangoes | Mango | Historic | Fruit | (Yadav & Singh, 2017) |
| *Musa* | *sp.* | Bananas | Muz | Historic | Fruit | (Neumann & Hildebrand, 2009; Nayar, 2010; Power *et al.*, 2019) |
| *Opuntia* | *ficus-indica* | Prickly pear | Qulqual | Late modern | Fruit | (Gebremeskel G *et al.*, 2013) |
| *Passiflora* | *edulis* | Passionfruit | Passionfruit | Late modern | Fruit | (Lakew *et al.*, 2018) |
| *Persea* | *americana* | Avocados | Abokado | Late modern | Fruit | (Berhanu, 2013) |
| *Phoenix* | *reclinata* | Date palm | Zenbaba | Indigenous | Fruit | (Plants of the World Online \| Kew Science 2021) |
| *Prunus* | *persica* | Peach | Kok | Historic | Fruit | (Goettsch, 1991) |
| *Psidium* | *guajava* | Guavas | Zeytun | Early modern | Fruit | (Harlan, 1969) |
| *Arachis* | *hypogaea* | Groundnuts | Lewz/Occoloni | Early modern | Oil | (Harlan, 1969) |
| *Brassica* | *nigra* | Black mustard | Senafc | Indigenous | Oil | (Goettsch, 1991) |
| *Brassica* | *napus* | Rapeseed | Gomen zer | Historic | Oil | (Harlan, 1969; Goettsch, 1991) |
| *Helianthus* | *annuus* | Sunflower | Suf | Early modern | Oil | (Harlan, 1969) |
| *Linum* | *usitatissimum* | Flax/Lin seed | Telba | Historic | Oil | (Harlan, 1969; Goettsch, 1991) |
| *Ricinus* | *communis* | Castor bean | Gulo | Indigenous | Oil | (Harlan, 1969; Goettsch, 1991) |
| *Sesamum* | *indicum* | Sesame | Selit | Historic | Oil | (Goettsch, 1991) |
| *Olea* | *eruopaea* | Olives | Weyra | Historic | Other | (Phillipson, 2012; Diez, 2015) |
| *Saccharum* | *officinarum* | Sugar cane | Shenkora ageda | Historic | Other | (Tena *et al.*, 2016; Power *et al.*, 2019) |
| *Cicer* | *arietinum* | Chickpea | Shmbra | Historic | Pulse | (Harlan, 1969; Phillipson, 2012) |
| *Glycine* | *max* | Soya | Akwri-ater | Late modern | Pulse | (Goettsch, 1991) |
| *Phaseolus* | *lunatus* | Lima beans | Adangware | Early modern | Pulse | (Harlan, 1969) |
| *Phaseolus* | *vulgaris* | Haricot beans | Bolqe | Early modern | Pulse | (Harlan, 1969; Goettsch, 1991) |
| *Pisum* | *sativum* | Peas | Ater | Historic | Pulse | (Harlan, 1969; Phillipson, 2012) |
| *Vicia* | *faba* | Faba/horse bean | baqela | Historic | Pulse | (Harlan, 1969; Phillipson, 2012) |
| *Allium* | *cepa* | Onion | Qey shinkurt | Historic | Root | (Harlan, 1969; Goettsch, 1991) |
| *Allium* | *sativum* | Garlic | Nec shinkurt | Early modern | Root | (Harlan, 1969; Goettsch, 1991) |
| *Beta* | *vulgaris* | Beet | Key sir | Late modern | Veg | (Harlan, 1969; Goettsch, 1991) |
| *Canna* | *indica* | Arrowroot | Seit akuri | Early modern | Root | (Gebrehiwot & Hundera, 2014; Asmelash, 2021) |
| *Colocasia* | *esculenta* | Taro | Goderie | Historic | Root | (Matthews, 2010; Power *et al.*, 2019) |
| *Daucus* | *carota* | Carrot | Karot | Late modern | Root | (Goettsch, 1991; Mohammed *et al.*, 2014) |
| *Dioscorea* | *bulbifera* | Arial yam | Hare kotea | Indigenous | Root | (Power *et al.*, 2019; ‘Plants of the World Online \| Kew Science’, 2021) |
| *Dioscorea* | *sp.* | Yam | Boye | Indigenous | Root | (Harlan, 1969; Goettsch, 1991; Power *et al.*, 2019) |
| *Ensete* | *ventricosum* | Enset | Enset/Koba | Indigenous | Root | (Harlan, 1969) |
| *Ipomoea* | *batatas* | Sweet potatoes | Siquar dinich | Early modern | Root | (Mohammed *et al.*, 2015) |
| *Manihot* | *esculenta* | Cassava | Yencet boye | Early modern | Root | (Okigbo, 1980) |
| *Plectranthus* | *edulis* | Ethiopian potato | Oromo dinich | Indigenous | Root | (Goettsch, 1991; Mekbib & Weibull, 2012) |
| *Solanum* | *tuberosum* | Potatoes | dinich | Late modern | Root | (Kolech *et al.*, 2015) |
| *Artemisia* | *afra/abyssinica* | Artemisia | Ch'uk'un/Arrity | Indigenous | Spice | (Plants of the World Online \| Kew Science 2021) |
| *Aframomum* | *corrorima* | Korarima | Korerima | Indigenous | Spice | (Jansen, 1981; ‘Plants of the World Online \| Kew Science’, 2021) |
| *Capsicum* | *anuum* | Chilli pepper | Qaraya | Early modern | Spice | (Harlan, 1969; Jansen, 1981) |
| *Coriandrum* | *sativum* | Coriander | Dimbilal | Historic | Spice | (Harlan, 1969) |
| *Cymbopogon* | *citratus* | lemongrass | Tej sar | Historic | Spice | (Jansen, 1981) |
| *Foeniculum* | *vulgare* | Fennel | Inselal | Late modern | Spice | (Goettsch, 1991) |
| *Lippia* | *abyssinica* | Koseret | Koseret | Indigenous | Spice | (Plants of the World Online \| Kew Science 2021) |
| *Ocimum* | *Americanum* | sacred basil | Besobilla | Indigenous | Spice | (Goettsch, 1991) |
| *Ocimum* | *Lamiifolium* | Basil | Damakesse | Indigenous | Spice | (Plants of the World Online \| Kew Science 2021) |
| *Rhamnus* | *prioides* | Hops | Gesho | Indigenous | Spice | (Harlan, 1969; Zeven & Zhukovsky, 1975) |
| *Salvia* | *Rosmarinus* | Rosemary | Yesiga metibesha | Historic | Spice | (Goettsch, 1991; ‘Plants of the World Online \| Kew Science’, 2021) |
| *Ruta* | *chalepenesis* | Fringed Rue | Tena adam | Historic | Spice | (Jansen, 1981) |
| *Satanocrater* | *somalensis* | Tea spice | Yeshai kimem | Indigenous | Spice | (Plants of the World Online \| Kew Science 2021) |
| *Thymus* | *schimperi* | Thyme | tosigne | Indigenous | Spice | (Plants of the World Online \| Kew Science 2021) |
| *Zingiber* | *officinale* | Ginger | zingible | Historic | Spice | (Harlan, 1969; Goettsch, 1991) |
| *Camellia* | *sinensis* | Tea | Chai kitel | Late modern | Stimulant | (Zakir, 2017) |
| *Catha* | *edulis* | Chat | Cat | Indigenous | Stimulant | (Harlan, 1969) |
| *Coffea* | *arabica* | Coffee | Buna | Indigenous | Stimulant | (Harlan, 1969; ‘Plants of the World Online \| Kew Science’, 2021) |
| *Nicotiana* | *sp.* | Tobacco | Tinbaho | Early modern | Stimulant | (Zeven & Zhukovsky, 1975) |
| *Beta* | *vulgaris* | Chard | Qosta | Late modern | Veg | (Harlan, 1969; Goettsch, 1991) |
| *Brassica* | *carinata* | Ethiopian cabbage | Gomen zer | Indigenous | Veg | (Goettsch, 1991) |
| *Brassica* | *oleracea* | Cabbage | Tql gomen | Historic | Veg | (Goettsch, 1991) |
| *Brassica* | *rapa* | Napa cabbage | Hamli | Historic | Veg | (Harlan 1969; Plants of the World Online \| Kew Science 2021) |
| *Commelina* | *africana* | Commelina | Yewef' nqur | Indigenous | Veg | (Plants of the World Online \| Kew Science 2021) |
| *Cucurbita* | *sp.* | Pumpkin/squash | Duba | Early modern | Veg | (Harlan, 1969) |
| *Lactuca* | *sativa* | Lettuce | Selata | Late modern | Veg | (Harlan, 1969) |
| *Lycopersicon* | *esculentum* | Tomatoes | Timatim | Early modern | Veg | (Harlan, 1969) |
| *Moringa* | *sp.* | Shiferaw | Shiferaw | Indigenous | Veg | (Goettsch, 1991; ‘Plants of the World Online \| Kew Science’, 2021) |
| *Solanum* | *nigrum* | - | Tikur awitt | Indigenous | Veg | (Plants of the World Online \| Kew Science 2021) |
| *Solanum* | *tarderemotum* | - |  | Indigenous | Veg | (Plants of the World Online \| Kew Science 2021) |

**Table S2.** Crop presence and cultivated area for 1,369 Ethiopian farms, with associated socioeconomic metadata.

*See separate file.*

### Supplementary Figures

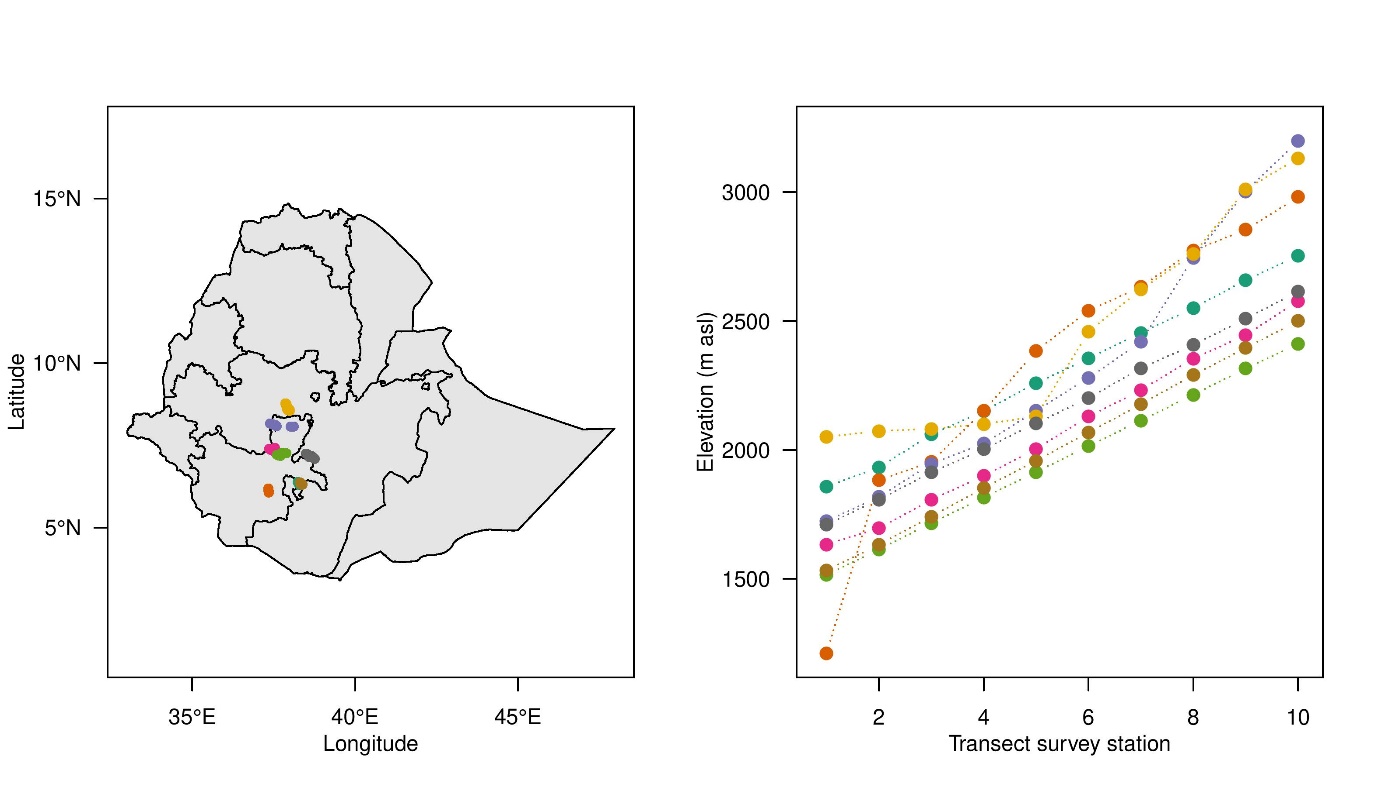

**Figure S1.** Characteristics of farm surveys conducted in southwestern Ethiopia, showing a) the location of eight distinct transects and b) the elevations where sampling took place. Points denote main sampling stations with 15-20 farms surveyed.

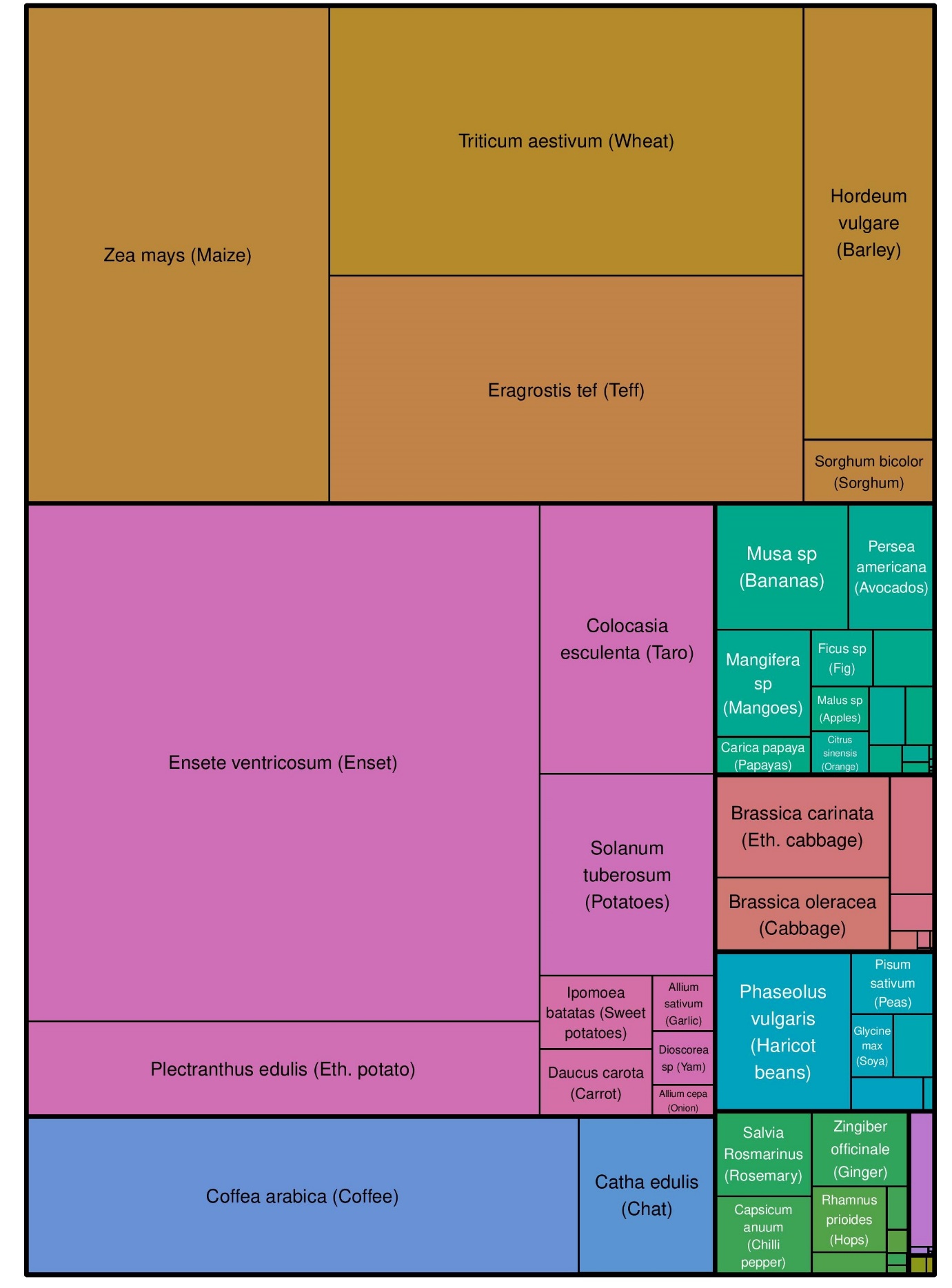

**Figure S2. Hierarchical treemap visualisation of crops in this study, with rectangle size proportional to area.** Colours denote crop types.

**USAID**. **2016a**. *Spatial Data Repository, The Demographic and Health Surveys Program. Modeled Surfaces. ET2016DHS_CHVACCCBAS_MS_v01 ICF International. Funded by the United States Agency for International Development (USAID). Available from spatialdata.dhsprogram.com. [Acc*.

**USAID**. **2016b**. *Spatial Data Repository, The Demographic and Health Surveys Program. Modeled Surfaces. ET2016DHS_CNNUTSCHA2_MS_v01 ICF International. Funded by the United States Agency for International Development (USAID). Available from spatialdata.dhsprogram.com. [Acc*.

**USAID**. **2016c**. : *Spatial Data Repository, The Demographic and Health Surveys Program. Modeled Surfaces. ET2016DHS_EDLITRMLIT_MS_v01 ICF International. Funded by the United States Agency for International Development (USAID). Available from spatialdata.dhsprogram.com. [A*.

**USAID**. **2016d**. : *Spatial Data Repository, The Demographic and Health Surveys Program. Modeled Surfaces. ET2016DHS_EDLITRWLIT_MS_v01 ICF International. Funded by the United States Agency for International Development (USAID). Available from spatialdata.dhsprogram.com. [A*.

**USAID**. **2016e**. *Spatial Data Repository, The Demographic and Health Surveys Program. Modeled Surfaces. ET2016DHS_FPNADMWUNT_MS_v01 ICF International. Funded by the United States Agency for International Development (USAID). Available from spatialdata.dhsprogram.com. [Acc*.

**USAID**. **2016f**. *Spatial Data Repository, The Demographic and Health Surveys Program. Modeled Surfaces. ET2016DHS_WSSRCEPIMP_MS_v01 ICF International. Funded by the United States Agency for International Development (USAID). Available from spatialdata.dhsprogram.com. [Acc*.

**Yadav D, Singh SP**. **2017**. Mango: History origin and distribution. *Journal of Pharmaconosy and Phytochemistry* **6**: 1257–1262.

**Zakir M**. **2017**. Review on Tea (Camellia sinensis) Research Achievements, Challenges and Future Prospective Including Ethiopian Status. *International Journal of Forestry and Horticulture* **3**: 27–39.

**Zeven AC, Zhukovsky PM**. **1975**. *Dictionary of cultivated plants and their centres of diversity*. Wageningen: Center for Agricultural Publishing and Documentation.
